## Supplementary material for "The interplay between temperature and growth phase shapes the transcriptional landscape of *Pseudomonas aeruginosa*": Table S1

**Table S1. Genes whose LasR regulation depends on temperature.**

| PA Locus | Gene Name | Gene Description | LasR Regulation at 37°C | LasR Regulation at 25°C | How Temperature Affects LasR Regulation |
|---|---|---|---|---|---|
| PA0024 | hemF | coproporphyrinogen III oxidase, aerobic | -0.019819667 | 1.289756578 | Negatively LasR regulated at 25°C |
| PA0025 | aroE | shikimate dehydrogenase | 0.052683089 | 1.12647734 | Negatively LasR regulated at 25°C |
| PA0029 | | probable sulfate transporter | -1.137766372 | 0.744816807 | Positively LasR regulated at 37°C |
| PA0038 | | hypothetical protein | -0.676269034 | -1.535485651 | Positively LasR regulated at 25°C |
| PA0043 | | hypothetical protein | -1.038102778 | -0.166703581 | Positively LasR regulated at 37°C |
| PA0047 | | hypothetical protein | 0.388587597 | 1.475127645 | Negatively LasR regulated at 25°C |
| PA0048 | | probable transcriptional regulator | -1.300288029 | -0.146092638 | Positively LasR regulated at 37°C |
| PA0049 | | hypothetical protein | -2.365962739 | -0.503480184 | Positively LasR regulated at 37°C |
| PA0060 | | conserved hypothetical protein | -0.306391201 | -1.103046006 | Positively LasR regulated at 25°C |
| PA0061 | | hypothetical protein | -1.156020018 | -0.434606109 | Positively LasR regulated at 37°C |
| PA0062 | | hypothetical protein | -1.037193126 | -0.977094655 | Positively LasR regulated at 37°C |
| PA0072 | tagS1 | TagS1 | -1.148772338 | -0.16202482 | Positively LasR regulated at 37°C |
| PA0086 | tagJ1 | TagJ1 | -0.14723237 | 1.109192444 | Negatively LasR regulated at 25°C |
| PA0097 | | hypothetical protein | -0.357175623 | 1.216855906 | Negatively LasR regulated at 25°C |
| PA0104 | | hypothetical protein | -1.504441707 | -0.323706045 | Positively LasR regulated at 37°C |

| | | | | | |
|---|---|---|---|---|---|
| PA0109 | | hypothetical protein | -1.456427165 | -0.443991218 | Positively LasR regulated at 37°C |
| PA0128 | | conserved hypothetical protein | 1.064971095 | -0.004728532 | Negatively LasR regulated at 37°C |
| PA0129 | bauD | Amino acid permease | 1.05608781 | -0.355484843 | Negatively LasR regulated at 37°C |
| PA0152 | pcaQ | transcriptional regulator PcaQ | -1.166602384 | -0.944500494 | Positively LasR regulated at 37°C |
| PA0159 | | probable transcriptional regulator | -0.520735942 | -1.335171723 | Positively LasR regulated at 25°C |
| PA0160 | | hypothetical protein | -0.17385085 | -1.238596058 | Positively LasR regulated at 25°C |
| PA0162 | opdC | histidine porin OpdC | -0.514417287 | -1.285037503 | Positively LasR regulated at 25°C |
| PA0169 | siaD | SiaD | 1.626711549 | -0.189970392 | Negatively LasR regulated at 37°C |
| PA0170 | siaC | SiaC | 1.301206233 | -0.436603986 | Negatively LasR regulated at 37°C |
| PA0171 | siaB | SiaB | 2.066780482 | -0.190019324 | Negatively LasR regulated at 37°C |
| PA0172 | siaA | SiaA | 1.450727072 | 0.219287244 | Negatively LasR regulated at 37°C |
| PA0177 | | probable purine-binding chemotaxis protein | -1.013597719 | -0.350385342 | Positively LasR regulated at 37°C |
| PA0189 | | probable porin | -0.99886364 | -2.119533557 | Positively LasR regulated at 25°C |
| PA0194 | | hypothetical protein | -1.528202006 | -0.431579288 | Positively LasR regulated at 37°C |
| PA0200 | | hypothetical protein | 0.118771696 | 1.164022796 | Negatively LasR regulated at 25°C |
| PA0208 | mdcA | malonate decarboxylase alpha subunit | 0.467838772 | 1.972280884 | Negatively LasR regulated at 25°C |
| PA0209 | | conserved hypothetical protein | -0.009600823 | 1.946643017 | Negatively LasR regulated at 25°C |

| | | | | | |
|---|---|---|---|---|---|
| PA0210 | mdcC | malonate decarboxylase delta subunit | 0.250520919 | 1.395039784 | Negatively LasR regulated at 25°C |
| PA0212 | mdcE | malonate decarboxylase gamma subunit | 0.763131141 | 1.758386077 | Negatively LasR regulated at 25°C |
| PA0213 | | hypothetical protein | 0.595059573 | 4.209656225 | Negatively LasR regulated at 25°C |
| PA0214 | | probable acyl transferase | 1.343049379 | 0.719958435 | Negatively LasR regulated at 37°C |
| PA0215 | | malonate transporter MadL | 1.624493971 | 0.595815457 | Negatively LasR regulated at 37°C |
| PA0230 | pcaB | 3-carboxy-cis,cis-muconate cycloisomerase | -1.129609742 | -0.682410609 | Positively LasR regulated at 37°C |
| PA0231 | pcaD | beta-ketoadipate enol-lactone hydrolase | -1.335007309 | -0.686009586 | Positively LasR regulated at 37°C |
| PA0263 | hcpC | secreted protein Hcp | 1.780251219 | -3.235073575 | Mixed LasR regulation between temperatures |
| PA0263.1 | | tRNA-Arg | 1.175923194 | 0.832378118 | Negatively LasR regulated at 37°C |
| PA0281 | cysW | sulfate transport protein CysW | 1.195487103 | 0.269859122 | Negatively LasR regulated at 37°C |
| PA0282 | cysT | sulfate transport protein CysT | 1.108979921 | 0.139228462 | Negatively LasR regulated at 37°C |
| PA0283 | sbp | sulfate-binding protein precursor | 1.671076583 | 0.543674134 | Negatively LasR regulated at 37°C |
| PA0284 | | hypothetical protein | 1.599445082 | 0.19907242 | Negatively LasR regulated at 37°C |
| PA0291 | oprE | Anaerobically-induced outer membrane porin OprE precursor | 1.087050415 | -0.230275983 | Negatively LasR regulated at 37°C |
| PA0298 | spuB | Glutamylpolyamine synthetase | 1.079068264 | 0.806205583 | Negatively LasR regulated at 37°C |

| | | | | | |
|---|---|---|---|---|---|
| PA0309 | | hypothetical protein | -1.014691995 | -0.92343533 | Positively LasR regulated at 37°C |
| PA0321 | | acetylpolyamine amidohydrolase | 0.1546683 | 1.295900319 | Negatively LasR regulated at 25°C |
| PA0354 | | conserved hypothetical protein | -0.83097435 | -1.564050799 | Positively LasR regulated at 25°C |
| PA0369 | | Uncharacterized protein | -0.664658431 | -1.301933785 | Positively LasR regulated at 25°C |
| PA0397 | | probable cation efflux system protein | -1.056826101 | -0.44483745 | Positively LasR regulated at 37°C |
| PA0423.1 | AS1974 | AS1974 | -1.300483465 | -0.50418423 | Positively LasR regulated at 37°C |
| PA0423.2 | AS1974-shor | AS1974-shorter1 | -1.298180501 | -0.512820973 | Positively LasR regulated at 37°C |
| PA0446 | | conserved hypothetical protein | -1.651459697 | 0.385544902 | Positively LasR regulated at 37°C |
| PA0447 | gcdH | glutaryl-CoA dehydrogenase | -1.221200928 | 0.255900437 | Positively LasR regulated at 37°C |
| PA0451 | | conserved hypothetical protein | -1.50586847 | 0.157458234 | Positively LasR regulated at 37°C |
| PA0467 | | conserved hypothetical protein | -1.04354138 | -0.385621683 | Positively LasR regulated at 37°C |
| PA0483 | | probable acetyltransferase | -1.343765998 | -0.471005616 | Positively LasR regulated at 37°C |
| PA0484 | | conserved hypothetical protein | -1.122952704 | -0.270385369 | Positively LasR regulated at 37°C |
| PA0485 | | conserved hypothetical protein | 1.282196545 | 0.806748091 | Negatively LasR regulated at 37°C |
| PA0509 | nirN | NirN | 0.407092482 | 4.550442098 | Negatively LasR regulated at 25°C |
| PA0510 | nirE | NirE | -0.460356279 | 4.820614432 | Negatively LasR regulated at 25°C |

| | | | | | |
|---|---|---|---|---|---|
| PA0511 | nirJ | heme d1 biosynthesis protein NirJ | 0.128630364 | 4.797635162 | Negatively LasR regulated at 25°C |
| PA0512 | nirH | NirH | 0.311098707 | 4.76874134 | Negatively LasR regulated at 25°C |
| PA0513 | nirG | NirG | -0.38856153 | 4.708172634 | Negatively LasR regulated at 25°C |
| PA0514 | nirL | heme d1 biosynthesis protein NirL | -0.860141254 | 4.75826201 | Negatively LasR regulated at 25°C |
| PA0515 | | probable transcriptional regulator | -0.302641251 | 4.761891378 | Negatively LasR regulated at 25°C |
| PA0516 | nirF | heme d1 biosynthesis protein NirF | -0.441403412 | 4.62772784 | Negatively LasR regulated at 25°C |
| PA0517 | nirC | probable c-type cytochrome precursor | -0.418694362 | 5.165982593 | Negatively LasR regulated at 25°C |
| PA0518 | nirM | cytochrome c-551 precursor | -0.474805967 | 5.371409507 | Negatively LasR regulated at 25°C |
| PA0519 | nirS | nitrite reductase precursor | -0.091209123 | 4.718615097 | Negatively LasR regulated at 25°C |
| PA0520 | nirQ | regulatory protein NirQ | -0.041969944 | 2.782596219 | Negatively LasR regulated at 25°C |
| PA0521 | | probable cytochrome c oxidase subunit | -0.635798204 | 4.341641027 | Negatively LasR regulated at 25°C |
| PA0522 | | hypothetical protein | -0.576464188 | 4.458203356 | Negatively LasR regulated at 25°C |
| PA0523 | norC | nitric-oxide reductase subunit C | 0.175033622 | 4.742116225 | Negatively LasR regulated at 25°C |
| PA0524 | norB | nitric-oxide reductase subunit B | -0.32238777 | 5.802947655 | Negatively LasR regulated at 25°C |

| | | | | | |
|---|---|---|---|---|---|
| PA0525 | | probable dinitrification protein NorD | -1.45944272 | 7.540251045 | Mixed LasR regulation between temperatures |
| PA0526 | | hypothetical protein | -1.114679122 | 2.693912679 | Mixed LasR regulation between temperatures |
| PA0543 | | hypothetical protein | -1.631785023 | -0.872430157 | Positively LasR regulated at 37°C |
| PA0554 | | hypothetical protein | 0.057453491 | -1.071656568 | Positively LasR regulated at 25°C |
| PA0574.1 | | tRNA-Met | 1.068158721 | 0.690898467 | Negatively LasR regulated at 37°C |
| PA0621 | | conserved hypothetical protein | 1.205172428 | 0.594467608 | Negatively LasR regulated at 37°C |
| PA0622 | | probable bacteriophage protein | 1.065386979 | 0.667245479 | Negatively LasR regulated at 37°C |
| PA0623 | | probable bacteriophage protein | 1.13880713 | 0.619261691 | Negatively LasR regulated at 37°C |
| PA0654 | speD | S-adenosylmethionine decarboxylase proenzyme | 1.761738637 | 0.299776517 | Negatively LasR regulated at 37°C |
| PA0670 | | hypothetical protein | -1.052062344 | -0.500758429 | Positively LasR regulated at 37°C |
| PA0673 | | hypothetical protein | -1.135082441 | 0.132514645 | Positively LasR regulated at 37°C |
| PA0676 | vreR | sigma factor regulator, VreR | 0.496310295 | 1.150669372 | Negatively LasR regulated at 25°C |
| PA0700 | | hypothetical protein | -1.748572366 | 1.250192781 | Mixed LasR regulation between temperatures |
| PA0704 | | probable amidase | -1.103732496 | -0.584512628 | Positively LasR regulated at 37°C |
| PA0709 | | hypothetical protein | -0.30880051 | -1.097300466 | Positively LasR regulated at 25°C |

| | | | | | |
|---|---|---|---|---|---|
| PA0710 | gloA2 | lactoylglutathione lyase | -0.23485019 | -2.276673277 | Positively LasR regulated at 25°C |
| PA0730 | | probable transferase | -0.019447703 | -1.197296091 | Positively LasR regulated at 25°C |
| PA0737 | | hypothetical protein | -0.476930799 | -1.127885958 | Positively LasR regulated at 25°C |
| PA0741 | | conserved hypothetical protein | 1.034380651 | 0.419860066 | Negatively LasR regulated at 37°C |
| PA0742 | | hypothetical protein | 0.093980334 | 1.058584287 | Negatively LasR regulated at 25°C |
| PA0743 | | probable 3-hydroxyisobutyrate dehydrogenase | -1.006723967 | -0.145681359 | Positively LasR regulated at 37°C |
| PA0751 | | conserved hypothetical protein | -1.294448927 | -0.36902798 | Positively LasR regulated at 37°C |
| PA0753 | | hypothetical protein | -1.825807273 | -0.29423146 | Positively LasR regulated at 37°C |
| PA0788 | | hypothetical protein | -1.395557895 | -0.307845289 | Positively LasR regulated at 37°C |
| PA0798 | pmtA | phospholipid methyltransferase | -1.019303708 | -0.529942925 | Positively LasR regulated at 37°C |
| PA0803 | | hypothetical protein | -1.10311954 | -0.127772881 | Positively LasR regulated at 37°C |
| PA0806 | | hypothetical protein | -1.427680114 | -0.115078297 | Positively LasR regulated at 37°C |
| PA0813 | | hypothetical protein | -1.632599684 | -0.550365402 | Positively LasR regulated at 37°C |
| PA0836.1 | P5 | P5 | 0.344939522 | 1.140003543 | Negatively LasR regulated at 25°C |
| PA0844 | plcH | hemolytic phospholipase C precursor | -1.018200894 | 0.303008061 | Positively LasR regulated at 37°C |

| | | | | | |
|---|---|---|---|---|---|
| PA0885 | | probable C4-dicarboxylate transporter | -1.131471776 | 0.905936395 | Positively LasR regulated at 37°C |
| PA0894 | | hypothetical protein | -1.251169103 | 0.679644249 | Positively LasR regulated at 37°C |
| PA0905.1 | | tRNA-Ser | 1.163476309 | 0.255822878 | Negatively LasR regulated at 37°C |
| PA0905.2 | | tRNA-Arg | 1.087224221 | 0.274732306 | Negatively LasR regulated at 37°C |
| PA0905.3 | | tRNA-Arg | 1.036134305 | 0.280706984 | Negatively LasR regulated at 37°C |
| PA0911 | alpE | AlpE | 1.343772725 | 0.625705634 | Negatively LasR regulated at 37°C |
| PA0947 | | conserved hypothetical protein | 1.116193102 | 0.003114435 | Negatively LasR regulated at 37°C |
| PA0961 | | probable cold-shock protein | 1.392340192 | 0.019692189 | Negatively LasR regulated at 37°C |
| PA0964 | pmpR | pqsR-mediated PQS regulator, PmpR | 1.20544007 | 0.280681115 | Negatively LasR regulated at 37°C |
| PA0972 | tolB | TolB protein | 1.052628787 | 0.401397622 | Negatively LasR regulated at 37°C |
| PA0975 | | probable radical activating enzyme | 1.453445433 | 0.519300642 | Negatively LasR regulated at 37°C |
| PA0976.1 | | tRNA-Lys | 1.379132251 | -0.033066316 | Negatively LasR regulated at 37°C |
| PA0980 | | hypothetical protein | -1.210746352 | -0.910074088 | Positively LasR regulated at 37°C |
| PA1013 | purC | phosphoribosylaminoimidazole-succinocarboxamide synthase | 1.190402366 | 0.269737774 | Negatively LasR regulated at 37°C |
| PA1034 | | hypothetical protein | 1.231262751 | 0.170366201 | Negatively LasR regulated at 37°C |
| PA1051 | | probable transporter | 0.891943961 | 1.204883984 | Negatively LasR regulated at 25°C |
| PA1070 | braG | branched-chain amino acid transport protein BraG | 0.400069438 | 1.125194635 | Negatively LasR regulated at 25°C |

| | | | | | |
|---|---|---|---|---|---|
| PA1071 | braF | branched-chain amino acid transport protein BraF | 0.710056073 | 1.326665408 | Negatively LasR regulated at 25°C |
| PA1072 | braE | branched-chain amino acid transport protein BraE | 0.37687369 | 1.386432219 | Negatively LasR regulated at 25°C |
| PA1073 | braD | branched-chain amino acid transport protein BraD | 0.990477288 | 1.175387695 | Negatively LasR regulated at 25°C |
| PA1074 | braC | branched-chain amino acid transport protein BraC | 0.815469057 | 1.176279845 | Negatively LasR regulated at 25°C |
| PA1075 | | hypothetical protein | 0.891150155 | 1.063382992 | Negatively LasR regulated at 25°C |
| PA1112.1 | | | -1.016348456 | -0.081104657 | Positively LasR regulated at 37°C |
| PA1114 | | hypothetical protein | -0.356513502 | -1.208559628 | Positively LasR regulated at 25°C |
| PA1116 | | hypothetical protein | 1.040897602 | 0.115559501 | Negatively LasR regulated at 37°C |
| PA1123 | | hypothetical protein | 1.483968727 | 0.775531735 | Negatively LasR regulated at 37°C |
| PA1147 | | probable amino acid permease | -0.188634829 | 1.231823513 | Negatively LasR regulated at 25°C |
| PA1166 | | hypothetical protein | -1.111010344 | -0.545419586 | Positively LasR regulated at 37°C |
| PA1176 | napF | ferredoxin protein NapF | -1.42004259 | -0.600365322 | Positively LasR regulated at 37°C |
| PA1177 | napE | periplasmic nitrate reductase protein NapE | -1.062805675 | -0.476566986 | Positively LasR regulated at 37°C |

| | | | | | |
|---|---|---|---|---|---|
| PA1195 | ddaH | dimethylarginine dimethylaminohydrolase DdaH | 0.433813382 | 1.221410675 | Negatively LasR regulated at 25°C |
| PA1212 | | probable major facilitator superfamily (MFS) transporter | -0.930621149 | -1.401838204 | Positively LasR regulated at 25°C |
| PA1225 | | FAD-dependent NADPH:quinone reductase | 1.527033982 | -1.674965287 | Mixed LasR regulation between temperatures |
| PA1252 | dpkA | DpkA | -1.012943033 | -0.712786495 | Positively LasR regulated at 37°C |
| PA1260 | lhpP | ABC transporter periplasmic-binding protein, LhpP | -0.080915131 | 1.864817085 | Negatively LasR regulated at 25°C |
| PA1268 | lhpA | Hydroxyproline 2-epimerase, LhpA | 1.037205021 | -2.949172374 | Mixed LasR regulation between temperatures |
| PA1282 | | probable major facilitator superfamily (MFS) transporter | 1.722170227 | 0.97513232 | Negatively LasR regulated at 37°C |
| PA1287 | | probable glutathione peroxidase | -0.99984826 | -1.273736381 | Positively LasR regulated at 25°C |
| PA1288 | odsT | oxylipin transporter | 0.71318577 | 1.055138356 | Negatively LasR regulated at 25°C |
| PA1300 | hxuI | HxuI | -0.549119949 | 1.921377649 | Negatively LasR regulated at 25°C |
| PA1312 | | probable transcriptional regulator | -1.182354873 | -0.376045703 | Positively LasR regulated at 37°C |
| PA1317 | cyoA | cytochrome o ubiquinol oxidase subunit II | 0.057830045 | -1.254178574 | Positively LasR regulated at 25°C |
| PA1318 | cyoB | cytochrome o ubiquinol oxidase subunit I | 0.96379718 | -1.552322496 | Positively LasR regulated at 25°C |

| | | | | | |
|---|---|---|---|---|---|
| PA1319 | cyoC | cytochrome o ubiquinol oxidase subunit III | -0.211818401 | -1.211585875 | Positively LasR regulated at 25°C |
| PA1324.1 | P9 | P9 | 1.080458761 | -0.222630155 | Negatively LasR regulated at 37°C |
| PA1327 | | probable protease | -1.255838496 | -0.455245902 | Positively LasR regulated at 37°C |
| PA1349 | | conserved hypothetical protein | -1.584890078 | -0.623115462 | Positively LasR regulated at 37°C |
| PA1351 | | probable sigma-70 factor, ECF subfamily | -1.262026242 | -0.009269003 | Positively LasR regulated at 37°C |
| PA1394 | | hypothetical protein | -0.04169931 | 1.668737418 | Negatively LasR regulated at 25°C |
| PA1407 | | hypothetical protein | -0.606761801 | -1.47916709 | Positively LasR regulated at 25°C |
| PA1410 | | probable periplasmic spermidine/putrescine-binding protein | -0.372827009 | 1.235995018 | Negatively LasR regulated at 25°C |
| PA1415 | | hypothetical protein | -1.209956307 | 0.078298151 | Positively LasR regulated at 37°C |
| PA1418 | | probable sodium:solute symport protein | 0.403360628 | 1.606916263 | Negatively LasR regulated at 25°C |
| PA1421 | gbuA | guanidinobutyrase | -0.398876464 | 2.24799619 | Negatively LasR regulated at 25°C |
| PA1425 | | probable ATP-binding component of ABC transporter | 1.217843434 | 0.205743914 | Negatively LasR regulated at 37°C |
| PA1428 | | conserved hypothetical protein | 1.03852653 | 0.581507985 | Negatively LasR regulated at 37°C |
| PA1471 | | hypothetical protein | -1.404252963 | -0.72260092 | Positively LasR regulated at 37°C |
| PA1485 | | probable amino acid permease | -0.173916389 | 1.271400701 | Negatively LasR regulated at 25°C |

| | | | | | |
|---|---|---|---|---|---|
| PA1486 | bapF | beta-peptidyl aminopeptidase | 0.533121669 | 1.255804201 | Negatively LasR regulated at 25°C |
| PA1488 | | hypothetical protein | -2.785286304 | 0.424903011 | Positively LasR regulated at 37°C |
| PA1504 | | probable transcriptional regulator | 1.074620279 | 0.187130663 | Negatively LasR regulated at 37°C |
| PA1511 | vgrG2a | VgrG2a | 0.142300963 | -1.439006868 | Positively LasR regulated at 25°C |
| PA1519 | | probable transporter | 0.166938767 | 1.479336197 | Negatively LasR regulated at 25°C |
| PA1580 | gltA | citrate synthase | 1.518968516 | 0.735887061 | Negatively LasR regulated at 37°C |
| PA1592 | | hypothetical protein | -1.17292989 | -0.970538494 | Positively LasR regulated at 37°C |
| PA1593 | | hypothetical protein | 1.017883324 | -0.042819485 | Negatively LasR regulated at 37°C |
| PA1599 | | probable transcriptional regulator | -1.184880935 | -0.892489934 | Positively LasR regulated at 37°C |
| PA1605 | | hypothetical protein | -0.857504579 | -1.395250226 | Positively LasR regulated at 25°C |
| PA1606 | | hypothetical protein | -0.6015786 | -1.52001954 | Positively LasR regulated at 25°C |
| PA1608 | | probable chemotaxis transducer | 0.75075681 | 1.584593061 | Negatively LasR regulated at 25°C |
| PA1620 | | hypothetical protein | 0.069161791 | 3.57782156 | Negatively LasR regulated at 25°C |
| PA1651 | | probable transporter | 1.069304705 | 0.521689464 | Negatively LasR regulated at 37°C |
| PA1673 | mhr | microoxic hemerythrin, Mhr | 0.818613498 | 1.603354007 | Negatively LasR regulated at 25°C |
| PA1687 | speE | spermidine synthase | 1.226371061 | -0.151303314 | Negatively LasR regulated at 37°C |

| | | | | | |
|---|---|---|---|---|---|
| PA1689 | | conserved hypothetical protein | 0.230705848 | -1.421979412 | Positively LasR regulated at 25°C |
| PA1733 | | conserved hypothetical protein | -1.000308083 | -0.40748751 | Positively LasR regulated at 37°C |
| PA1740 | | hypothetical protein | -1.154590972 | 1.111303789 | Mixed LasR regulation between temperatures |
| PA1745 | | hypothetical protein | -1.010798363 | -0.839867981 | Positively LasR regulated at 37°C |
| PA1747 | | hypothetical protein | 0.493936311 | 1.152239729 | Negatively LasR regulated at 25°C |
| PA1757 | thrH | homoserine kinase | 1.40834578 | 0.173206679 | Negatively LasR regulated at 37°C |
| PA1786 | nasS | NasS | -1.209343512 | -0.319643316 | Positively LasR regulated at 37°C |
| PA1793 | ppiB | peptidyl-prolyl cis-trans isomerase B | 1.113442982 | 0.294860253 | Negatively LasR regulated at 37°C |
| PA1796.2 | | tRNA-His | 1.021230151 | 0.509397707 | Negatively LasR regulated at 37°C |
| PA1796.3 | | tRNA-Leu | 1.136648506 | 0.601700436 | Negatively LasR regulated at 37°C |
| PA1796.4 | | tRNA-His | 1.011007248 | 0.700362453 | Negatively LasR regulated at 37°C |
| PA1826 | | probable transcriptional regulator | -1.135994337 | -0.202308941 | Positively LasR regulated at 37°C |
| PA1830 | | hypothetical protein | 1.021036051 | 0.195453008 | Negatively LasR regulated at 37°C |
| PA1831 | | hypothetical protein | 1.101545117 | 0.050420811 | Negatively LasR regulated at 37°C |
| PA1835 | | hypothetical protein | -1.114074637 | -0.549299318 | Positively LasR regulated at 37°C |
| PA1868 | xqhA | secretion protein XqhA | -1.554466229 | -0.53062911 | Positively LasR regulated at 37°C |
| PA1880 | | probable oxidoreductase | -1.161258062 | -0.361847815 | Positively LasR regulated at 37°C |
| PA1898 | qscR | quorum-sensing control repressor | 0.132164013 | -1.836671467 | Positively LasR regulated at 25°C |

| | | | | | |
|---|---|---|---|---|---|
| PA1911 | femR | sigma factor regulator, FemR | 0.15626453 | 2.293392108 | Negatively LasR regulated at 25°C |
| PA1912 | femI | ECF sigma factor, FemI | 0.980560656 | 1.263660294 | Negatively LasR regulated at 25°C |
| PA1939 | | hypothetical protein | -0.486542911 | -1.172809298 | Positively LasR regulated at 25°C |
| PA1948 | rbsC | membrane protein component of ABC ribose transporter | 0.261417222 | 1.06885216 | Negatively LasR regulated at 25°C |
| PA1973 | pqqF | pyrroloquinoline quinone biosynthesis protein F | 1.05407308 | 0.789714983 | Negatively LasR regulated at 37°C |
| PA1980 | eraR | response regulator EraR | -0.185985656 | 2.109406981 | Negatively LasR regulated at 25°C |
| PA1997 | | probable AMP-binding enzyme | 0.469373994 | 1.403987988 | Negatively LasR regulated at 25°C |
| PA2024 | | probable ring-cleaving dioxygenase | -1.166256868 | -0.293371436 | Positively LasR regulated at 37°C |
| PA2038 | | hypothetical protein | 1.361700076 | 0.164815854 | Negatively LasR regulated at 37°C |
| PA2063 | | hypothetical protein | 1.298060267 | 0.999164431 | Negatively LasR regulated at 37°C |
| PA2074 | | hypothetical protein | -0.928221133 | -1.893650379 | Positively LasR regulated at 25°C |
| PA2075 | | hypothetical protein | -1.028123642 | -0.432963584 | Positively LasR regulated at 37°C |
| PA2085 | | probable ring-hydroxylating dioxygenase small subunit | -1.03150813 | 2.757594385 | Mixed LasR regulation between temperatures |
| PA2109 | | hypothetical protein | 0.858143361 | 1.127671832 | Negatively LasR regulated at 25°C |

| | | | | | |
|---|---|---|---|---|---|
| PA2110 | | hypothetical protein | 0.506027239 | 1.117905807 | Negatively LasR regulated at 25°C |
| PA2111 | | hypothetical protein | 0.600656407 | 1.068959671 | Negatively LasR regulated at 25°C |
| PA2113 | opdO | pyroglutamate porin OpdO | 0.939359527 | 1.241176932 | Negatively LasR regulated at 25°C |
| PA2114 | | probable major facilitator superfamily (MFS) transporter | 0.954155658 | 1.138617288 | Negatively LasR regulated at 25°C |
| PA2115 | | probable transcriptional regulator | 1.335894652 | 0.322143273 | Negatively LasR regulated at 37°C |
| PA2128 | cupA1 | fimbrial subunit CupA1 | 1.210042065 | 0.154697232 | Negatively LasR regulated at 37°C |
| PA2129 | cupA2 | chaperone CupA2 | 1.500349893 | -0.055591874 | Negatively LasR regulated at 37°C |
| PA2130 | cupA3 | usher CupA3 | 1.274057171 | -0.112061684 | Negatively LasR regulated at 37°C |
| PA2131 | cupA4 | fimbrial subunit CupA4 | 1.394381698 | 0.491261463 | Negatively LasR regulated at 37°C |
| PA2177 | | probable sensor/response regulator hybrid | -1.10209519 | -0.818113981 | Positively LasR regulated at 37°C |
| PA2181 | | hypothetical protein | -1.896550706 | -0.934257742 | Positively LasR regulated at 37°C |
| PA2188 | | probable alcohol dehydrogenase (Zn-dependent) | -0.871513477 | -1.690393604 | Positively LasR regulated at 25°C |
| PA2189 | | hypothetical protein | -0.978250081 | -1.239318038 | Positively LasR regulated at 25°C |
| PA2196 | | TetR family transcriptional regulator | -0.933383032 | -1.243515849 | Positively LasR regulated at 25°C |
| PA2197 | | conserved hypothetical protein | -0.775919884 | -1.119501154 | Positively LasR regulated at 25°C |

| | | | | | |
|---|---|---|---|---|---|
| PA2202 | | probable amino acid permease | 2.122612913 | 0.563888662 | Negatively LasR regulated at 37°C |
| PA2203 | | probable amino acid permease | 1.996235142 | 0.537080719 | Negatively LasR regulated at 37°C |
| PA2236 | pslF | PslF | -1.056044293 | -0.501415509 | Positively LasR regulated at 37°C |
| PA2241 | pslK | PslL | -1.225626255 | -0.064778756 | Positively LasR regulated at 37°C |
| PA2301 | | hypothetical protein | -3.273248089 | -0.906453815 | Positively LasR regulated at 37°C |
| PA2306 | ambA | AmbA | 0.145510188 | -1.025923341 | Positively LasR regulated at 25°C |
| PA2319 | | probable transposase | 1.225647495 | 0.962107915 | Negatively LasR regulated at 37°C |
| PA2338 | | probable binding protein component of ABC maltose/mannitol transporter | 0.474229853 | 1.027699639 | Negatively LasR regulated at 25°C |
| PA2339 | | probable binding-protein-dependent maltose/mannitol transport protein | 0.72239883 | 2.740106835 | Negatively LasR regulated at 25°C |
| PA2340 | | probable binding-protein-dependent maltose/mannitol transport protein | 0.226820502 | 1.705918613 | Negatively LasR regulated at 25°C |
| PA2341 | | probable ATP-binding component of ABC maltose/mannitol transporter | 0.575242818 | 1.921612993 | Negatively LasR regulated at 25°C |
| PA2348 | | conserved hypothetical protein | -1.746317546 | 3.746137905 | Mixed LasR regulation between temperatures |
| PA2351 | | probable permease of ABC transporter | -1.404353642 | 0.637921958 | Positively LasR regulated at 37°C |

| | | | | | |
|---|---|---|---|---|---|
| PA2374 | tseF | TseF | -1.010534819 | -0.50060406 | Positively LasR regulated at 37°C |
| PA2375 | | hypothetical protein | -1.219996747 | -0.753918173 | Positively LasR regulated at 37°C |
| PA2378 | | probable aldehyde dehydrogenase | -1.037782587 | -0.336846282 | Positively LasR regulated at 37°C |
| PA2384 | | hypothetical protein | -2.192872324 | -0.852252461 | Positively LasR regulated at 37°C |
| PA2424 | pvdL | PvdL | -3.275762416 | -0.552329691 | Positively LasR regulated at 37°C |
| PA2431 | | hypothetical protein | -1.596843467 | -0.354176274 | Positively LasR regulated at 37°C |
| PA2435 | | probable cation-transporting P-type ATPase | -1.149211379 | -0.39405128 | Positively LasR regulated at 37°C |
| PA2448 | | putative hydrolase | -2.282029602 | -0.939964576 | Positively LasR regulated at 37°C |
| PA2451 | | hypothetical protein | -2.900096404 | 0.14010675 | Positively LasR regulated at 37°C |
| PA2465 | | hypothetical protein | 0.576368877 | -1.541272398 | Positively LasR regulated at 25°C |
| PA2537 | | probable acyltransferase | 0.853210567 | 1.822501508 | Negatively LasR regulated at 25°C |
| PA2538 | | hypothetical protein | 0.680917636 | 1.295034002 | Negatively LasR regulated at 25°C |
| PA2539 | | conserved hypothetical protein | 0.709238817 | 2.302422216 | Negatively LasR regulated at 25°C |
| PA2541 | | probable CDP-alcohol phosphatidyltransferase | 0.79951196 | 1.778568455 | Negatively LasR regulated at 25°C |
| PA2559.1 | srfA | SrfA | -1.155547169 | -0.90968203 | Positively LasR regulated at 37°C |
| PA2567 | | hypothetical protein | 0.60246774 | 1.029002582 | Negatively LasR regulated at 25°C |
| PA2569 | | hypothetical protein | -1.118625035 | -0.762406555 | Positively LasR regulated at 37°C |

| | | | | | |
|---|---|---|---|---|---|
| PA2572 | | probable two-component response regulator | -1.197229748 | -0.374634805 | Positively LasR regulated at 37°C |
| PA2575 | | hypothetical protein | 1.258021455 | 0.885687262 | Negatively LasR regulated at 37°C |
| PA2619 | infA | initiation factor | 1.170529913 | -0.02676956 | Negatively LasR regulated at 37°C |
| PA2629 | purB | adenylosuccinate lyase | 1.053892268 | 0.108002406 | Negatively LasR regulated at 37°C |
| PA2636 | | hypothetical protein | 0.563066331 | 2.743607421 | Negatively LasR regulated at 25°C |
| PA2637 | nuoA | NADH dehydrogenase I chain A | 1.560416603 | 0.262809525 | Negatively LasR regulated at 37°C |
| PA2662 | | conserved hypothetical protein | -0.297702077 | 1.602008748 | Negatively LasR regulated at 25°C |
| PA2664 | fhp | flavohemoprotein | -0.992835793 | -2.058437049 | Positively LasR regulated at 25°C |
| PA2697 | | hypothetical protein | -1.270706791 | -0.72095389 | Positively LasR regulated at 37°C |
| PA2701 | | probable major facilitator superfamily (MFS) transporter | -1.220814095 | -0.470899677 | Positively LasR regulated at 37°C |
| PA2708 | | hypothetical protein | -0.736814205 | -1.145313455 | Positively LasR regulated at 25°C |
| PA2763 | | hypothetical protein | -2.427284217 | 0.489523983 | Positively LasR regulated at 37°C |
| PA2787 | cpg2 | carboxypeptidase G2 precursor | -1.015480889 | -0.627685125 | Positively LasR regulated at 37°C |
| PA2789 | | hypothetical protein | 1.133252773 | 0.423491599 | Negatively LasR regulated at 37°C |
| PA2799 | | hypothetical protein | -1.176408834 | -0.305078552 | Positively LasR regulated at 37°C |
| PA2819.1 | | tRNA-Gly | 1.441757513 | 0.300209226 | Negatively LasR regulated at 37°C |
| PA2819.2 | | tRNA-Gly | 1.0856697 | 0.51755994 | Negatively LasR regulated at 37°C |

| | | | | | |
|---|---|---|---|---|---|
| PA2819.3 | | tRNA-Glu | 1.062010987 | 0.512724166 | Negatively LasR regulated at 37°C |
| PA2828 | | probable aminotransferase | 1.010014169 | 0.09741274 | Negatively LasR regulated at 37°C |
| PA2829 | | hypothetical protein | 1.738110922 | 0.18432867 | Negatively LasR regulated at 37°C |
| PA2831 | | conserved hypothetical protein | -0.647280062 | -1.222677435 | Positively LasR regulated at 25°C |
| PA2835 | | probable major facilitator superfamily (MFS) transporter | -1.617518624 | 0.394632725 | Positively LasR regulated at 37°C |
| PA2847 | | conserved hypothetical protein | -1.157151623 | -0.218275097 | Positively LasR regulated at 37°C |
| PA2851 | efp | translation elongation factor P | 1.242782817 | 0.006481752 | Negatively LasR regulated at 37°C |
| PA2862 | lipA | lactonizing lipase precursor | 0.560945176 | 1.380524574 | Negatively LasR regulated at 25°C |
| PA2911 | | probable TonB-dependent receptor | 0.552741981 | 1.211552061 | Negatively LasR regulated at 25°C |
| PA2920 | | probable chemotaxis transducer | -1.02961526 | -0.721139216 | Positively LasR regulated at 37°C |
| PA2984 | | hypothetical protein | -1.073236613 | -0.256123017 | Positively LasR regulated at 37°C |
| PA2991 | sth | soluble pyridine nucleotide transhydrogenase | 1.134102371 | 0.583683742 | Negatively LasR regulated at 37°C |
| PA3009 | | hypothetical protein | 1.172081335 | 0.556905203 | Negatively LasR regulated at 37°C |
| PA3041 | | hypothetical protein | -1.013871513 | -0.918046844 | Positively LasR regulated at 37°C |
| PA3060 | pelE | PelE | -1.007340607 | 0 | Positively LasR regulated at 37°C |

| | | | | | |
|---|---|---|---|---|---|
| PA3063 | pelB | PelB | -0.882602669 | -1.026593956 | Positively LasR regulated at 25°C |
| PA3089 | | hypothetical protein | -1.297629441 | -0.489735647 | Positively LasR regulated at 37°C |
| PA3133.1 | | tRNA-Glu | 1.060494554 | 0.685670681 | Negatively LasR regulated at 37°C |
| PA3141 | wbpM | nucleotide sugar epimerase/dehydratase WbpM | -0.352048898 | -1.078816281 | Positively LasR regulated at 25°C |
| PA3149 | wbpH | probable glycosyltransferase WbpH | -0.356187271 | -1.054341216 | Positively LasR regulated at 25°C |
| PA3162 | rpsA | 30S ribosomal protein S1 | 1.408741592 | -0.102794016 | Negatively LasR regulated at 37°C |
| PA3183 | zwf | glucose-6-phosphate 1-dehydrogenase | 1.013167691 | 0.472881971 | Negatively LasR regulated at 37°C |
| PA3209 | | conserved hypothetical protein | 0.751331485 | 1.247230342 | Negatively LasR regulated at 25°C |
| PA3249 | | probable transcriptional regulator | -0.453866334 | -1.288925437 | Positively LasR regulated at 25°C |
| PA3262.1 | | tRNA-Asp | 1.672914424 | 0.808354855 | Negatively LasR regulated at 37°C |
| PA3262.2 | | tRNA-Val | 1.636615209 | 0.799122468 | Negatively LasR regulated at 37°C |
| PA3269 | | probable transcriptional regulator | 1.026866902 | 0.300608519 | Negatively LasR regulated at 37°C |
| PA3291 | tli1 | Tli1 | 0.132333916 | -1.214889784 | Positively LasR regulated at 25°C |
| PA3292 | | hypothetical protein | 0.015644339 | -1.2179584 | Positively LasR regulated at 25°C |
| PA3311 | nbdA | NbdA | -1.195949594 | -0.132193764 | Positively LasR regulated at 37°C |
| PA3315 | | probable permease of ABC transporter | -2.34777539 | -0.209156799 | Positively LasR regulated at 37°C |

| | | | | | |
|---|---|---|---|---|---|
| PA3316 | | probable permease of ABC transporter | -1.475642371 | 0.134542706 | Positively LasR regulated at 37°C |
| PA3340 | | hypothetical protein | -1.231088185 | 0.043985048 | Positively LasR regulated at 37°C |
| PA3360 | | probable secretion protein | 1.287486687 | 0.754744454 | Negatively LasR regulated at 37°C |
| PA3366.1 | amiL | AmiL | 1.636139678 | 0.88052104 | Negatively LasR regulated at 37°C |
| PA3369 | | hypothetical protein | -0.820468033 | -1.454999073 | Positively LasR regulated at 25°C |
| PA3391 | nosR | regulatory protein NosR | -0.906802925 | 2.728094525 | Negatively LasR regulated at 25°C |
| PA3392 | nosZ | nitrous-oxide reductase precursor | 0.464047815 | 6.594431901 | Negatively LasR regulated at 25°C |
| PA3393 | nosD | NosD protein | -0.221230951 | 6.604069507 | Negatively LasR regulated at 25°C |
| PA3394 | nosF | NosF protein | 0.553595986 | 6.870319031 | Negatively LasR regulated at 25°C |
| PA3395 | nosY | NosY protein | -1.035814114 | 6.960704444 | Mixed LasR regulation between temperatures |
| PA3396 | nosL | NosL protein | -4.069500523 | 6.207716233 | Mixed LasR regulation between temperatures |
| PA3407 | hasAp | heme acquisition protein HasAp | -1.990712864 | 1.119004337 | Mixed LasR regulation between temperatures |
| PA3409 | hasS | HasS | 0.392565691 | 1.719449437 | Negatively LasR regulated at 25°C |
| PA3410 | hasI | HasI | 0.467186188 | 1.620011269 | Negatively LasR regulated at 25°C |
| PA3416 | | probable pyruvate dehydrogenase E1 component, beta chain | -1.334178049 | -0.468278518 | Positively LasR regulated at 37°C |
| PA3417 | | probable pyruvate dehydrogenase E1 component, alpha subunit | -1.331606328 | -0.006873809 | Positively LasR regulated at 37°C |
| PA3418 | ldh | leucine dehydrogenase | -1.040394381 | -0.315377746 | Positively LasR regulated at 37°C |

| | | | | | |
|---|---|---|---|---|---|
| PA3426 | | probable enoyl CoA-hydratase/isomerase | -1.035209882 | -0.530639183 | Positively LasR regulated at 37°C |
| PA3431 | | conserved hypothetical protein | 0.262037579 | 1.293801939 | Negatively LasR regulated at 25°C |
| PA3432 | | hypothetical protein | 0.106372187 | 1.327242026 | Negatively LasR regulated at 25°C |
| PA3434 | | probable transposase | 1.644032638 | 0.765616829 | Negatively LasR regulated at 37°C |
| PA3441 | | probable molybdopterin-binding protein | 0.807185782 | 1.269235659 | Negatively LasR regulated at 25°C |
| PA3447 | | probable ATP-binding component of ABC transporter | -1.319741697 | 2.539267309 | Mixed LasR regulation between temperatures |
| PA3450 | lsfA | 1-Cys peroxiredoxin LsfA | 1.635089473 | 0.438532437 | Negatively LasR regulated at 37°C |
| PA3457 | | hypothetical protein | -0.531440963 | -1.088269085 | Positively LasR regulated at 25°C |
| PA3475 | pheC | cyclohexadienyl dehydratase precursor | -1.771039155 | -0.719836517 | Positively LasR regulated at 37°C |
| PA3486 | vgrG4b | VgrG4b | 0.047930129 | -2.928305037 | Positively LasR regulated at 25°C |
| PA3487 | tle5 | Tle5 | -0.332792139 | -2.093287525 | Positively LasR regulated at 25°C |
| PA3488 | tli5 | Tli5 | -0.017995788 | -2.078247675 | Positively LasR regulated at 25°C |
| PA3500 | | conserved hypothetical protein | 0.071991141 | 1.017365052 | Negatively LasR regulated at 25°C |
| PA3501 | | hypothetical protein | -0.162721036 | 1.141142133 | Negatively LasR regulated at 25°C |
| PA3503 | | hypothetical protein | -0.917389402 | 1.508640032 | Negatively LasR regulated at 25°C |

| | | | | | |
|---|---|---|---|---|---|
| PA3510 | | hypothetical protein | 0.757446117 | 1.150768191 | Negatively LasR regulated at 25°C |
| PA3516 | | probable lyase | -0.803913445 | -1.692635684 | Positively LasR regulated at 25°C |
| PA3535.1 | realL | ReaL | 1.769220436 | 0.044380639 | Negatively LasR regulated at 37°C |
| PA3568 | | probable acetyl-coa synthetase | 0.648341008 | 1.341292231 | Negatively LasR regulated at 25°C |
| PA3585 | glpM | membrane protein GlpM | 1.134940149 | 0.433256818 | Negatively LasR regulated at 37°C |
| PA3598 | | conserved hypothetical protein | -0.86933794 | -1.397244353 | Positively LasR regulated at 25°C |
| PA3621.1 | rsmZ | regulatory RNA RsmZ | 1.114473105 | 0.620689552 | Negatively LasR regulated at 37°C |
| PA3632 | | conserved hypothetical protein | 1.065136037 | 0.115334679 | Negatively LasR regulated at 37°C |
| PA3641 | | probable amino acid permease | 1.2800681 | 0.468850216 | Negatively LasR regulated at 37°C |
| PA3655 | tsf | elongation factor Ts | 1.02169396 | -0.102637187 | Negatively LasR regulated at 37°C |
| PA3656 | rpsB | 30S ribosomal protein S2 | 1.357704708 | -0.100657962 | Negatively LasR regulated at 37°C |
| PA3700 | lysS | lysyl-tRNA synthetase | 1.236577742 | -0.034441634 | Negatively LasR regulated at 37°C |
| PA3709 | | probable major facilitator superfamily (MFS) transporter | 0.670015738 | 1.728170205 | Negatively LasR regulated at 25°C |
| PA3718 | | probable major facilitator superfamily (MFS) transporter | 0.626112786 | 1.077848172 | Negatively LasR regulated at 25°C |
| PA3720 | | hypothetical protein | 0.270112065 | 1.652531323 | Negatively LasR regulated at 25°C |

| | | | | | |
|---|---|---|---|---|---|
| PA3723 | | probable FMN oxidoreductase | -1.159074493 | -0.704040669 | Positively LasR regulated at 37°C |
| PA3729 | | conserved hypothetical protein | 1.039764983 | 0.739020987 | Negatively LasR regulated at 37°C |
| PA3739 | | probable sodium/hydrogen antiporter | -1.056294474 | -0.773074245 | Positively LasR regulated at 37°C |
| PA3741 | | hypothetical protein | 1.33860704 | 0.499567899 | Negatively LasR regulated at 37°C |
| PA3745 | rpsP | 30S ribosomal protein S16 | 1.016325062 | 0.164443645 | Negatively LasR regulated at 37°C |
| PA3757 | nagR | Transcriptional regulator of N-Acetylglucosamine catabolism operon | 0.755072962 | 1.054135052 | Negatively LasR regulated at 25°C |
| PA3766 | | probable aromatic amino acid transporter | 1.586445284 | 0.717050467 | Negatively LasR regulated at 37°C |
| PA3820 | secF | secretion protein SecF | 1.124397017 | -0.042767114 | Negatively LasR regulated at 37°C |
| PA3821 | secD | secretion protein SecD | 1.147748954 | -0.089180344 | Negatively LasR regulated at 37°C |
| PA3822 | | conserved hypothetical protein | 1.015162924 | -0.04986583 | Negatively LasR regulated at 37°C |
| PA3846 | | hypothetical protein | -1.285106295 | -0.436764868 | Positively LasR regulated at 37°C |
| PA3858 | | probable amino acid-binding protein | -1.356405285 | -0.12818939 | Positively LasR regulated at 37°C |
| PA3866 | | Pyocin S4 | 1.011278232 | 0.332250525 | Negatively LasR regulated at 37°C |
| PA3870 | moaA1 | molybdopterin biosynthetic protein A1 | 1.640481624 | 0.762459283 | Negatively LasR regulated at 37°C |

| | | | | | |
|---|---|---|---|---|---|
| PA3874 | narH | respiratory nitrate reductase beta chain | 0.571976976 | 1.690417805 | Negatively LasR regulated at 25°C |
| PA3877 | narK1 | nitrite extrusion protein 1 | 0.817577888 | 3.545840533 | Negatively LasR regulated at 25°C |
| PA3879 | narL | two-component response regulator NarL | 0.183874162 | 1.021421958 | Negatively LasR regulated at 25°C |
| PA3880 | | conserved hypothetical protein | 0.370218098 | 1.398625543 | Negatively LasR regulated at 25°C |
| PA3884 | | hypothetical protein | 0.631233753 | 1.007362275 | Negatively LasR regulated at 25°C |
| PA3893 | | conserved hypothetical protein | 0.115797197 | -1.51667075 | Positively LasR regulated at 25°C |
| PA3910 | eddA | Extracelullar DNA degradation protein, EddA | 0.316608989 | 2.045308428 | Negatively LasR regulated at 25°C |
| PA3911 | | conserved hypothetical protein | -0.09331023 | 2.633512625 | Negatively LasR regulated at 25°C |
| PA3912 | | conserved hypothetical protein | 0.075192864 | 2.556271084 | Negatively LasR regulated at 25°C |
| PA3913 | | probable protease | 0.224229438 | 2.163455205 | Negatively LasR regulated at 25°C |
| PA3920 | copA1 | CopA1 | -0.830673814 | -1.224711955 | Positively LasR regulated at 25°C |
| PA3925 | | probable acyl-CoA thiolase | 0.285499049 | 1.009784154 | Negatively LasR regulated at 25°C |
| PA3953 | | conserved hypothetical protein | 1.278970273 | 0.870699195 | Negatively LasR regulated at 37°C |
| PA3957 | | probable short-chain dehydrogenase | -1.129987332 | -0.293239792 | Positively LasR regulated at 37°C |

| | | | | | |
|---|---|---|---|---|---|
| PA3962 | | hypothetical protein | -0.560954784 | -1.137370444 | Positively LasR regulated at 25°C |
| PA3986 | | hypothetical protein | -1.069251275 | -0.351127784 | Positively LasR regulated at 37°C |
| PA3993 | | probable transposase | 1.064457034 | -0.551327481 | Negatively LasR regulated at 37°C |
| PA4005 | | conserved hypothetical protein | 1.041964986 | -0.037424636 | Negatively LasR regulated at 37°C |
| PA4017 | | thioesterase | -1.179726607 | -0.269899641 | Positively LasR regulated at 37°C |
| PA4023 | eat | ethanolamine transporter, Eat | 0.879440072 | 1.760000588 | Negatively LasR regulated at 25°C |
| PA4039 | | hypothetical protein | -0.807393133 | -1.44931209 | Positively LasR regulated at 25°C |
| PA4041 | | hypothetical protein | -1.4170441 | -0.382618945 | Positively LasR regulated at 37°C |
| PA4058 | | hypothetical protein | -1.251201967 | 0.106328094 | Positively LasR regulated at 37°C |
| PA4067 | oprG | Outer membrane protein OprG precursor | 0.623879536 | 1.219582428 | Negatively LasR regulated at 25°C |
| PA4072 | | probable amino acid permease | 0.838473322 | 1.012772638 | Negatively LasR regulated at 25°C |
| PA4073 | | probable aldehyde dehydrogenase | 0.718008134 | 1.18209742 | Negatively LasR regulated at 25°C |
| PA4090 | | hypothetical protein | 1.239107401 | 0.48216484 | Negatively LasR regulated at 37°C |
| PA4091 | hpaA | 4-hydroxyphenylacetate 3-monooxygenase large chain | 0.975614038 | 1.07751941 | Negatively LasR regulated at 25°C |

| | | | | | |
|---|---|---|---|---|---|
| PA4092 | hpaC | 4-hydroxyphenylacetate 3-monooxygenase small chain | 0.796746486 | 1.139062583 | Negatively LasR regulated at 25°C |
| PA4109 | ampR | transcriptional regulator AmpR | -1.285542022 | -0.256446989 | Positively LasR regulated at 37°C |
| PA4112 | | probable sensor/response regulator hybrid | -1.062624189 | -0.4936559 | Positively LasR regulated at 37°C |
| PA4127 | hpcG | 2-oxo-hept-3-ene-1,7-dioate hydratase | 1.796571608 | 0.654077768 | Negatively LasR regulated at 37°C |
| PA4148 | | probable short-chain dehydrogenase | -1.048864834 | 1.320639693 | Mixed LasR regulation between temperatures |
| PA4149 | | conserved hypothetical protein | 0.379863303 | 2.303441421 | Negatively LasR regulated at 25°C |
| PA4150 | | probable dehydrogenase E1 component | -0.34865464 | 1.669948717 | Negatively LasR regulated at 25°C |
| PA4152 | | probable hydrolase | 0.330911704 | 1.702897262 | Negatively LasR regulated at 25°C |
| PA4153 | | 2,3-butanediol dehydrogenase | -0.486088789 | 1.650588943 | Negatively LasR regulated at 25°C |
| PA4173 | | conserved hypothetical protein | -0.516074946 | -3.148038017 | Positively LasR regulated at 25°C |
| PA4194 | | probable permease of ABC transporter | -1.583779314 | 0.903049939 | Positively LasR regulated at 37°C |

| | | | | | |
|---|---|---|---|---|---|
| PA4206 | mexH | probable Resistance-Nodulation-Cell Division (RND) efflux membrane fusion protein precursor | -1.422151852 | -0.261822748 | Positively LasR regulated at 37°C |
| PA4208 | opmD | probable outer membrane protein precursor | -1.529049781 | 2.228974342 | Mixed LasR regulation between temperatures |
| PA4235 | ftnA | bacterial ferritin | 0.465922413 | 1.10452429 | Negatively LasR regulated at 25°C |
| PA4238 | rpoA | DNA-directed RNA polymerase alpha chain | 1.107792417 | -0.053419343 | Negatively LasR regulated at 37°C |
| PA4239 | rpsD | 30S ribosomal protein S4 | 1.183215408 | 0.096441818 | Negatively LasR regulated at 37°C |
| PA4245 | rpmD | 50S ribosomal protein L30 | 1.004837408 | -0.156669265 | Negatively LasR regulated at 37°C |
| PA4246 | rpsE | 30S ribosomal protein S5 | 1.253918565 | -0.121845478 | Negatively LasR regulated at 37°C |
| PA4247 | rplR | 50S ribosomal protein L18 | 1.096897178 | -0.268187487 | Negatively LasR regulated at 37°C |
| PA4261 | rplW | 50S ribosomal protein L23 | 1.139759267 | -0.158835334 | Negatively LasR regulated at 37°C |
| PA4263 | rplC | 50S ribosomal protein L3 | 1.076893831 | -0.12169116 | Negatively LasR regulated at 37°C |
| PA4268 | rpsL | 30S ribosomal protein S12 | 1.04432772 | -0.117495684 | Negatively LasR regulated at 37°C |
| PA4270.1 | P26 | P26 | 1.30157107 | -0.534495165 | Negatively LasR regulated at 37°C |
| PA4271 | rplL | 50S ribosomal protein L7 / L12 | 1.121201665 | -0.546521044 | Negatively LasR regulated at 37°C |
| PA4272 | rplJ | 50S ribosomal protein L10 | 1.289027192 | -0.454158586 | Negatively LasR regulated at 37°C |
| PA4273 | rplA | 50S ribosomal protein L1 | 1.131723453 | -0.221368024 | Negatively LasR regulated at 37°C |

| | | | | | |
|---|---|---|---|---|---|
| PA4274 | rplK | 50S ribosomal protein L11 | 1.139916615 | -0.151302769 | Negatively LasR regulated at 37°C |
| PA4289 | | probable transporter | -1.223503748 | -0.191861712 | Positively LasR regulated at 37°C |
| PA4292 | | probable phosphate transporter | 1.003798804 | 0.225397297 | Negatively LasR regulated at 37°C |
| PA4293 | pprA | two-component sensor PprA | -1.531574217 | -0.747887558 | Positively LasR regulated at 37°C |
| PA4294 | | hypothetical protein | -2.102823105 | -0.867063021 | Positively LasR regulated at 37°C |
| PA4296 | pprB | two-component response regulator, PprB | -1.079709378 | -0.568863731 | Positively LasR regulated at 37°C |
| PA4298 | | hypothetical protein | -0.764656762 | -2.685346428 | Positively LasR regulated at 25°C |
| PA4299 | tadD | TadD | -0.617413387 | -1.299909421 | Positively LasR regulated at 25°C |
| PA4300 | tadC | TadC | -0.794264133 | -1.171870148 | Positively LasR regulated at 25°C |
| PA4301 | tadB | TadB | -1.016241323 | -0.859610612 | Positively LasR regulated at 37°C |
| PA4305 | rcpC | RcpC | -1.560696149 | -0.686274969 | Positively LasR regulated at 37°C |
| PA4313 | | hypothetical protein | -1.162105913 | -0.328122086 | Positively LasR regulated at 37°C |
| PA4343 | | probable major facilitator superfamily (MFS) transporter | -1.251925702 | 0.047811163 | Positively LasR regulated at 37°C |
| PA4352 | | conserved hypothetical protein | -1.095760644 | 0.034273449 | Positively LasR regulated at 37°C |
| PA4354 | | conserved hypothetical protein | 1.003159241 | 0.462261997 | Negatively LasR regulated at 37°C |
| PA4362 | | hypothetical protein | -1.160062716 | -0.156843363 | Positively LasR regulated at 37°C |

| | | | | | |
|---|---|---|---|---|---|
| PA4375 | mexW | Resistance-Nodulation-Cell Division (RND) multidrug efflux transporter MexW | 1.075459516 | 0.291077258 | Negatively LasR regulated at 37°C |
| PA4394 | | conserved hypothetical protein | -0.284752947 | -1.138878855 | Positively LasR regulated at 25°C |
| PA4397 | panE | ketopantoate reductase | -1.203246853 | -0.203270686 | Positively LasR regulated at 37°C |
| PA4429 | | probable cytochrome c1 precursor | 1.010116883 | 0.285437582 | Negatively LasR regulated at 37°C |
| PA4430 | | probable cytochrome b | 1.014576056 | 0.262219321 | Negatively LasR regulated at 37°C |
| PA4432 | rpsI | 30S ribosomal protein S9 | 1.461107798 | 0.320262014 | Negatively LasR regulated at 37°C |
| PA4433 | rplM | 50S ribosomal protein L13 | 1.466841993 | 0.361614216 | Negatively LasR regulated at 37°C |
| PA4438 | | conserved hypothetical protein | 1.43757587 | 0.129135655 | Negatively LasR regulated at 37°C |
| PA4443 | cysD | ATP sulfurylase small subunit | 1.262039777 | 0.211864901 | Negatively LasR regulated at 37°C |
| PA4467 | | hypothetical protein | -1.970210497 | 1.365872343 | Mixed LasR regulation between temperatures |
| PA4494 | roxS | RoxS | 1.046739443 | 0.227873158 | Negatively LasR regulated at 37°C |
| PA4501 | opdD | Glycine-glutamate dipeptide porin OpdP | 0.165359097 | 2.163712808 | Negatively LasR regulated at 25°C |
| PA4502 | dppA4 | probable binding protein component of ABC transporter | -0.18431642 | 1.116053402 | Negatively LasR regulated at 25°C |

| | | | | | |
|---|---|---|---|---|---|
| PA4514 | | probable outer membrane receptor for iron transport | 0.988211681 | 2.246157987 | Negatively LasR regulated at 25°C |
| PA4516 | | hypothetical protein | -0.117101203 | 1.128438644 | Negatively LasR regulated at 25°C |
| PA4517 | | conserved hypothetical protein | 1.031501244 | 0.612674615 | Negatively LasR regulated at 37°C |
| PA4519 | speC | ornithine decarboxylase | -0.329783224 | -1.08997688 | Positively LasR regulated at 25°C |
| PA4535 | | hypothetical protein | -1.109313431 | -0.475934647 | Positively LasR regulated at 37°C |
| PA4541.3 | | tRNA-Asn | 0.998191816 | 1.013148627 | Negatively LasR regulated at 25°C |
| PA4563 | rpsT | 30S ribosomal protein S20 | 1.352617226 | 0.12327454 | Negatively LasR regulated at 37°C |
| PA4567 | rpmA | 50S ribosomal protein L27 | 1.476666405 | 0.058033644 | Negatively LasR regulated at 37°C |
| PA4568 | rplU | 50S ribosomal protein L21 | 1.406048641 | 0.003787696 | Negatively LasR regulated at 37°C |
| PA4573 | | hypothetical protein | -1.308488042 | -0.409089318 | Positively LasR regulated at 37°C |
| PA4574 | | conserved hypothetical protein | 1.672754679 | -0.081738277 | Negatively LasR regulated at 37°C |
| PA4575 | | hypothetical protein | -1.005844952 | -0.483725657 | Positively LasR regulated at 37°C |
| PA4583 | | conserved hypothetical protein | 1.132408588 | -0.049968933 | Negatively LasR regulated at 37°C |
| PA4589 | | probable outer membrane protein precursor | -0.880133278 | -1.907400094 | Positively LasR regulated at 25°C |
| PA4608 | mapZ | MapZ | -1.016753352 | -0.300803479 | Positively LasR regulated at 37°C |

| | | | | | |
|---|---|---|---|---|---|
| PA4610 | | hypothetical protein | -0.126426759 | 1.274155748 | Negatively LasR regulated at 25°C |
| PA4619 | | probable c-type cytochrome | 1.050540943 | 0.53596326 | Negatively LasR regulated at 37°C |
| PA4620 | | hypothetical protein | 0.907921053 | 1.003446323 | Negatively LasR regulated at 25°C |
| PA4621 | | probable oxidoreductase | 0.377366363 | 1.20133575 | Negatively LasR regulated at 25°C |
| PA4628 | lysP | lysine-specific permease | 1.009467811 | -0.121368192 | Negatively LasR regulated at 37°C |
| PA4629 | | hypothetical protein | 1.731597576 | -0.063366859 | Negatively LasR regulated at 37°C |
| PA4638 | | hypothetical protein | 1.104227895 | -0.805488304 | Negatively LasR regulated at 37°C |
| PA4640 | mqoB | malate:quinone oxidoreductase | 1.179719455 | 0.717823702 | Negatively LasR regulated at 37°C |
| PA4664 | prmC | S-adenosylmethionine-dependent methyltransferase, PrmC | 1.188631322 | 0.188373785 | Negatively LasR regulated at 37°C |
| PA4665 | prfA | peptide chain release factor 1 | 1.220916785 | 0.1279265 | Negatively LasR regulated at 37°C |
| PA4669.1 | | tRNA-Gln | 1.22683763 | 0.296370919 | Negatively LasR regulated at 37°C |
| PA4671 | | probable ribosomal protein L25 | 1.591429306 | -0.38713196 | Negatively LasR regulated at 37°C |
| PA4672 | | peptidyl-tRNA hydrolase | 1.287566134 | 0.008052326 | Negatively LasR regulated at 37°C |
| PA4673 | | conserved hypothetical protein | 1.577863348 | 0.208127474 | Negatively LasR regulated at 37°C |
| PA4676 | psCA3 | beta-carbonic anhydrase | -1.692578882 | -0.952354204 | Positively LasR regulated at 37°C |
| PA4685 | | hypothetical protein | 1.48654919 | -0.181568714 | Negatively LasR regulated at 37°C |

| | | | | | |
|---|---|---|---|---|---|
| PA4702 | | hypothetical protein | -1.267687546 | -0.612138338 | Positively LasR regulated at 37°C |
| PA4703 | | hypothetical protein | -1.879262012 | -0.785893926 | Positively LasR regulated at 37°C |
| PA4723 | | suppressor protein DksA | 1.157688325 | -0.062144124 | Negatively LasR regulated at 37°C |
| PA4726.2 | P30 | P30 | 1.423424668 | 0.71521447 | Negatively LasR regulated at 37°C |
| PA4766 | | conserved hypothetical protein | -1.243376848 | -0.377222953 | Positively LasR regulated at 37°C |
| PA4776 | pmrA | PmrA: two-component regulator system response regulator PmrA | -1.172740538 | -0.298975902 | Positively LasR regulated at 37°C |
| PA4780 | | conserved hypothetical protein | -1.018540206 | -0.253867362 | Positively LasR regulated at 37°C |
| PA4785 | | probable acyl-CoA thiolase | -0.660749131 | -1.276338987 | Positively LasR regulated at 25°C |
| PA4786 | | probable short-chain dehydrogenase | -0.594246399 | -1.095830582 | Positively LasR regulated at 25°C |
| PA4787 | | probable transcriptional regulator | -1.118983167 | 0.338349764 | Positively LasR regulated at 37°C |
| PA4788 | | hypothetical protein | -1.311157635 | -0.914172924 | Positively LasR regulated at 37°C |
| PA4802.1 | | tRNA-Sec | 1.386818133 | 0.339404105 | Negatively LasR regulated at 37°C |
| PA4843 | gcbA | GcbA | 0.792229605 | 1.182465638 | Negatively LasR regulated at 25°C |
| PA4853 | fis | DNA-binding protein Fis | 1.40737364 | 0.223665848 | Negatively LasR regulated at 37°C |
| PA4895 | | probable transmembrane sensor | -0.051793409 | 2.256584966 | Negatively LasR regulated at 25°C |

| | | | | | |
|---|---|---|---|---|---|
| PA4912 | | branched chain amino acid ABC transporter membrane protein | 1.05062993 | 0.511593525 | Negatively LasR regulated at 37°C |
| PA4916 | nrtR | Nudix-related transcriptional regulator NrtR | -3.138824421 | -0.900665377 | Positively LasR regulated at 37°C |
| PA4917 | nadD2 | nicotinate mononucleotide adenylyltransferase NadD2 | -2.85785432 | -0.843337334 | Positively LasR regulated at 37°C |
| PA4918 | pcnA | nicotinamidase, PcnA | 1.325977439 | 0.899312745 | Negatively LasR regulated at 37°C |
| PA4935 | rpsF | 30S ribosomal protein S6 | 1.024446734 | -0.183436873 | Negatively LasR regulated at 37°C |
| PA4937.1 | | tRNA-Leu | 1.147420491 | 0.604690895 | Negatively LasR regulated at 37°C |
| PA4974 | | probable outer membrane protein precursor | -0.628273046 | -1.159190911 | Positively LasR regulated at 25°C |
| PA5027 | | hypothetical protein | -1.761579659 | 0.092201549 | Positively LasR regulated at 37°C |
| PA5057 | phaD | poly(3-hydroxyalkanoic acid) depolymerase | -1.487372044 | -0.971287437 | Positively LasR regulated at 37°C |
| PA5076 | | putative amino acid ABC transporter substrate-binding protein | 1.099174844 | 0.243241467 | Negatively LasR regulated at 37°C |
| PA5083 | dguB | Rid2 subfamily protein | 0.508582444 | 1.110672077 | Negatively LasR regulated at 25°C |
| PA5084 | dguA | DguA | 1.359841619 | 0.724105703 | Negatively LasR regulated at 37°C |
| PA5087 | tli5b2 | type VI secretion lipase immunity protein, Tli5b2 | -0.142645293 | -1.074584453 | Positively LasR regulated at 25°C |

| | | | | | |
|---|---|---|---|---|---|
| PA5088 | tli5b3 | type VI secretion lipase immunity protein, Tli5b3 | 0.107903703 | -1.005763063 | Positively LasR regulated at 25°C |
| PA5089 | tle5b | type VI secretion phospholipase D effector Tle5b | 0.081337894 | -1.043097108 | Positively LasR regulated at 25°C |
| PA5096 | | probable binding protein component of ABC transporter | -0.722519542 | 1.054034838 | Negatively LasR regulated at 25°C |
| PA5097 | | probable amino acid permease | -0.224083981 | 1.287891667 | Negatively LasR regulated at 25°C |
| PA5098 | hutH | histidine ammonia-lyase | -0.36094981 | 1.193582374 | Negatively LasR regulated at 25°C |
| PA5137 | | hypothetical protein | 1.046525526 | 0.303597743 | Negatively LasR regulated at 37°C |
| PA5138 | | hypothetical protein | 1.100009427 | 0.438155765 | Negatively LasR regulated at 37°C |
| PA5153 | | amino acid (lysine/arginine/ornithine/histidine/octopine) ABC transporter periplasmic binding protein | 1.661624455 | 0.907236452 | Negatively LasR regulated at 37°C |
| PA5154 | | probable permease of ABC transporter | 1.750549199 | 0.69267987 | Negatively LasR regulated at 37°C |
| PA5155 | | amino acid (lysine/arginine/ornithine/histidine/octopine) ABC transporter membrane protein | 1.074195638 | 0.713888312 | Negatively LasR regulated at 37°C |

| | | | | | |
|---|---|---|---|---|---|
| PA5162 | rmlD | dTDP-4-dehydrorhamnose reductase | -1.447304091 | -0.940220256 | Positively LasR regulated at 37°C |
| PA5164 | rmlC | dTDP-4-dehydrorhamnose 3,5-epimerase | -1.01705357 | -0.684019304 | Positively LasR regulated at 37°C |
| PA5181 | | probable oxidoreductase | -0.501255247 | -1.456020329 | Positively LasR regulated at 25°C |
| PA5194 | | hypothetical protein | 1.026589361 | 0.411702838 | Negatively LasR regulated at 37°C |
| PA5205 | | conserved hypothetical protein | -0.442287249 | -2.151021189 | Positively LasR regulated at 25°C |
| PA5213 | gcvP1 | glycine cleavage system protein P1 | -1.078935199 | -0.427093754 | Positively LasR regulated at 37°C |
| PA5264 | | hypothetical protein | -0.101888735 | -1.427192276 | Positively LasR regulated at 25°C |
| PA5265 | | hypothetical protein | -0.175928181 | -1.595504974 | Positively LasR regulated at 25°C |
| PA5266 | vgrG6 | VgrG6 | -0.391023311 | -3.362498492 | Positively LasR regulated at 25°C |
| PA5267 | hcpB | secreted protein Hcp | 1.075311769 | -2.355044709 | Mixed LasR regulation between temperatures |
| PA5275 | | conserved hypothetical protein | 0.07807935 | 1.007419388 | Negatively LasR regulated at 25°C |
| PA5290 | | conserved hypothetical protein | -0.224497112 | -1.259648887 | Positively LasR regulated at 25°C |
| PA5291 | betT2 | BetT2 | -0.617225919 | -1.066646776 | Positively LasR regulated at 25°C |
| PA5295 | proE | ProE | -0.919460343 | -1.022283933 | Positively LasR regulated at 25°C |
| PA5298 | | xanthine phosphoribosyltransferase | 1.04021154 | -0.122660668 | Negatively LasR regulated at 37°C |

| | | | | | |
|---|---|---|---|---|---|
| PA5339 | | conserved hypothetical protein | 1.073928201 | 0.195825828 | Negatively LasR regulated at 37°C |
| PA5348 | | probable DNA-binding protein | 1.072471446 | 0.630686909 | Negatively LasR regulated at 37°C |
| PA5351 | rubA1 | Rubredoxin 1 | -0.444163188 | 1.252122449 | Negatively LasR regulated at 25°C |
| PA5352 | | conserved hypothetical protein | 0.119283048 | 1.619936937 | Negatively LasR regulated at 25°C |
| PA5353 | glcF | glycolate oxidase subunit GlcF | -0.37735687 | 1.407576281 | Negatively LasR regulated at 25°C |
| PA5354 | glcE | glycolate oxidase subunit GlcE | -0.307296786 | 1.2981524 | Negatively LasR regulated at 25°C |
| PA5367 | pstA | membrane protein component of ABC phosphate transporter | 1.132100776 | 0.601831485 | Negatively LasR regulated at 37°C |
| PA5368 | pstC | membrane protein component of ABC phosphate transporter | 1.624483886 | 0.747358152 | Negatively LasR regulated at 37°C |
| PA5394 | cls | cardiolipin synthase | -0.692784634 | -1.131624504 | Positively LasR regulated at 25°C |
| PA5403 | | probable transcriptional regulator | 1.284734039 | 0.275412299 | Negatively LasR regulated at 37°C |
| PA5408 | | hypothetical protein | -1.529225008 | 0.036443216 | Positively LasR regulated at 37°C |
| PA5409 | | hypothetical protein | -1.107891915 | -0.151226144 | Positively LasR regulated at 37°C |
| PA5423 | | hypothetical protein | -1.279028282 | -0.740150275 | Positively LasR regulated at 37°C |

| | | | | | |
|---|---|---|---|---|---|
| PA5426 | purE | phosphoribosylaminoimidazole carboxylase, catalytic subunit | 1.212568335 | -0.091305567 | Negatively LasR regulated at 37°C |
| PA5437 | | probable transcriptional regulator | 1.269660291 | 0.300512901 | Negatively LasR regulated at 37°C |
| PA5470 | | probable peptide chain release factor | -0.42363604 | 1.688147097 | Negatively LasR regulated at 25°C |
| PA5473 | | conserved hypothetical protein | -1.259936854 | -0.844852642 | Positively LasR regulated at 37°C |
| PA5476 | citA | citrate transporter | -0.64111253 | -1.100084988 | Positively LasR regulated at 25°C |
| PA5479 | gltP | proton-glutamate symporter | 1.151528323 | 0.357669757 | Negatively LasR regulated at 37°C |
| PA5483 | algB | two-component response regulator AlgB | -0.864741365 | -1.039269153 | Positively LasR regulated at 25°C |
| PA5484 | kinB | KinB | -0.918119686 | -1.162317964 | Positively LasR regulated at 25°C |
| PA5504 | | D-methionine ABC transporter membrane protein | 1.273285791 | 0.713155942 | Negatively LasR regulated at 37°C |
| PA5505 | | probable TonB-dependent receptor | 1.149850461 | 0.33097613 | Negatively LasR regulated at 37°C |
| PA5538 | amiA | N-acetylmuramoyl-L-alanine amidase | -0.130892384 | -1.439129637 | Positively LasR regulated at 25°C |
| PA5544 | | conserved hypothetical protein | 0.383663151 | 1.042804601 | Negatively LasR regulated at 25°C |
| PA5568 | | conserved hypothetical protein | 1.328945443 | 0.00947819 | Negatively LasR regulated at 37°C |

| | | | | | |
|---|---|---|---|---|---|
| PA5569 | rnpA | ribonuclease P protein component | 1.019380067 | 0.108734025 | Negatively LasR regulated at 37°C |
| PA5570 | rpmH | 50S ribosomal protein L34 | 1.168660646 | 0.012810537 | Negatively LasR regulated at 37°C |
